# Design and assessment of species-level qPCR primers targeting comammox

**DOI:** 10.1101/348664

**Authors:** Natalie Keene-Beach, Daniel R. Noguera

**Author notes:** Corresponding author: Daniel R. Noguera, 1415 Engineering Drive, Madison, WI 53706. E-mail addresses.

## Abstract

Published PCR primers targeting the ammonia monooxygenase gene (*amoA*) were applied to samples from activated sludge systems operated with low dissolved oxygen (DO) to quantify total and clade-level *Nitrospira* that perform complete ammonium oxidation (comammox); however, we found these existing primers resulted in significant artifact-associated non-target amplification. This not only overestimated comammox *amoA* copies but also resulted in numerous false positive detections in the environmental samples tested, as confirmed by gel electrophoresis. Therefore, to more accurately quantify known comammox, we designed specific and sensitive primers targeting three candidate species: *Candidatus* (*Ca.*) Nitrospira nitrosa, *Ca.* N. inopinata, and *Ca.* N. nitrificans. The primers were tested with *amoA* templates of these candidate species, and used to quantify comammox at the species level in low DO activated sludge systems. We found that comammox related to *Ca.* N. nitrosa were present and abundant in the majority of samples from low DO bioreactors and were not detected in samples from a high DO system. In addition, the greatest abundance of *Ca.* N. nitrosa was found in bioreactors operated with a long solids retention time. *Ca.* N. inopinata and *Ca.* N. nitrificans were only detected sporadically in these samples, indicating a minor role of these comammox in nitrification under low DO conditions.

**Abstract art.**
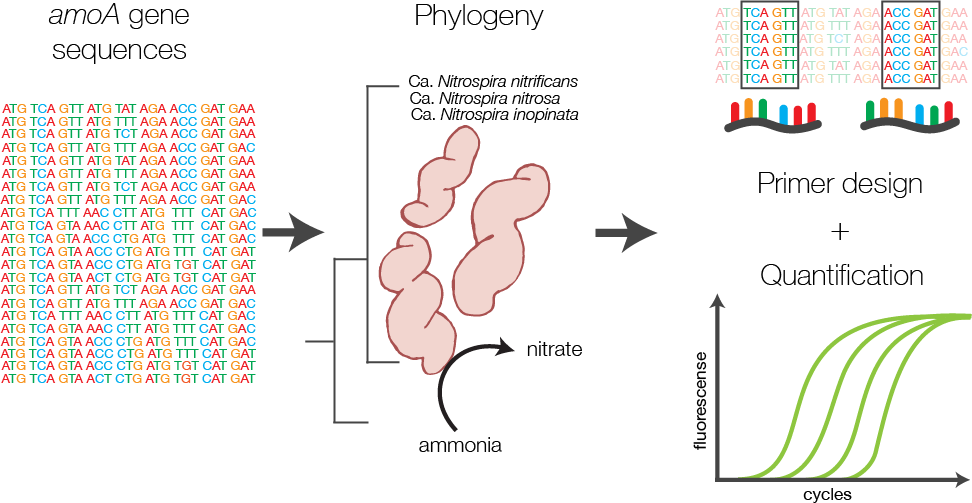

## Introduction

The oxidation of ammonium via nitrite to nitrate (i.e., nitrification) was historically considered a two-step process completed by phylogenetically distinct ammonia oxidizers and nitrite oxidizers. Although the complete oxidation of ammonium to nitrate by a single organism was theoretically possible, no microorganism with the ability to carry out both steps had been identified until late 2015.^1, 2^ These complete ammonium oxidizing (comammox) bacteria belong to the genus *Nitrospira*, which was known to contain only nitrite-oxidizing bacteria (NOB); therefore, an entire group of microorganisms with the ability to oxidize ammonia had been disguised for years.^1, 2^ The discovery of comammox bacteria has redefined a key component of the global nitrogen cycle, initiating a renewed focus of recent nitrogen cycling research in environmental biotechnology.^3–8^

Within the wastewater treatment industry, the application of nitrification under very low dissolved oxygen (DO) concentrations (below 0.2 mg O_2_/L) is an exciting energy-saving approach to biological nutrient removal (BNR)^9–11^. Previous attempts to identify the key microorganisms responsible for nitrification in low DO bioreactors have been inconclusive, with the presence of known ammonia oxidizing bacteria (AOB) and ammonia oxidizing archaea (AOA) unable to explain observed nitrification rates.^11^ Thus, the recent discovery of comammox in a variety of environments^1, 2^ prompted us to investigate their presence in low DO BNR systems.

The gene encoding the alpha-subunit of ammonia monooxygenase (*amoA*) has been one of the most widely used markers for detection and quantification of AOB and AOA, as it facilitates functional analysis and reconstruction of phylogenetic relationships.^12–15^ The *amoA* of comammox is distinguishable from *amoA* sequences of AOB and AOA, and thus, it can be used to detect the presence of comammox in environmental samples.^1–3, 16^ Recent descriptions of PCR primers targeting the *amoA* gene of commamox include a primer pair designed specifically for *Candidatus* (*Ca.*) Nitrospira inopinata,^1^ a primer pair designed specifically for a comammox-like clone within a freshwater aquaculture system,^5^ a primer collection that differentiates two broad clades of comammox within the *Nitrospira* genus,^17^ and a highly degenerate primer pair attempting to encompass all comammox within *Nitrospira*.^7^ To date, three candidate comammox species have been described in the literature, namely *Ca.* N. nitrosa, *Ca.* N. inopinata, and *Ca.* N. nitrificans.^1, 2^ However, quantifying the contribution of these candidate species to the comammox community in environmental samples has not been possible because of the lack of specific qPCR primer sets. To overcome this limitation, we designed a set of highly specific non-degenerate primers for the independent detection of *Ca.* N. nitrosa, *Ca.* N. inopinata, and *Ca.* N. nitrificans. We used these primers to evaluate the contribution of these species to the comammox population in samples from BNR plants operated at low DO conditions.

## Materials and Methods

### Sample collection, processing, and DNA extraction

Environmental samples used in this study originated from four low DO nitrifying bioreactors: a laboratory-scale sequencing batch reactor (L_SBR), a pilot-scale sequencing batch reactor (P_SBR), a pilot scale continuous flow (P_CF) reactor simulating a University of Cape Town configuration without nitrate recycle, and a full-scale wastewater treatment plant (WWTP) from the Trinity River Authority (TRA) Central Region (Arlington, TX) (Table 1). For comparison, samples were also collected from the full-scale Nine Springs WWTP (NS) at the Madison Metropolitan Sewerage District (Madison, WI), which operates with typical high DO conditions (Table 1). Biomass samples were stored at −80°C until DNA extraction. DNA was extracted using DNeasy® PowerSoil® DNA Isolation Kit (Qiagen, Hilden, Germany) following the manufacturer’s directions. DNA was quantified with a Qubit fluorometer (Thermo Fisher Scientific, Waltham, MA) and the purity ratio determined with a NanoDrop spectrophotometer (Thermo Fisher Scientific, Waltham, MA). DNA samples were stored at −20°C until further processing.

**Table 1.**
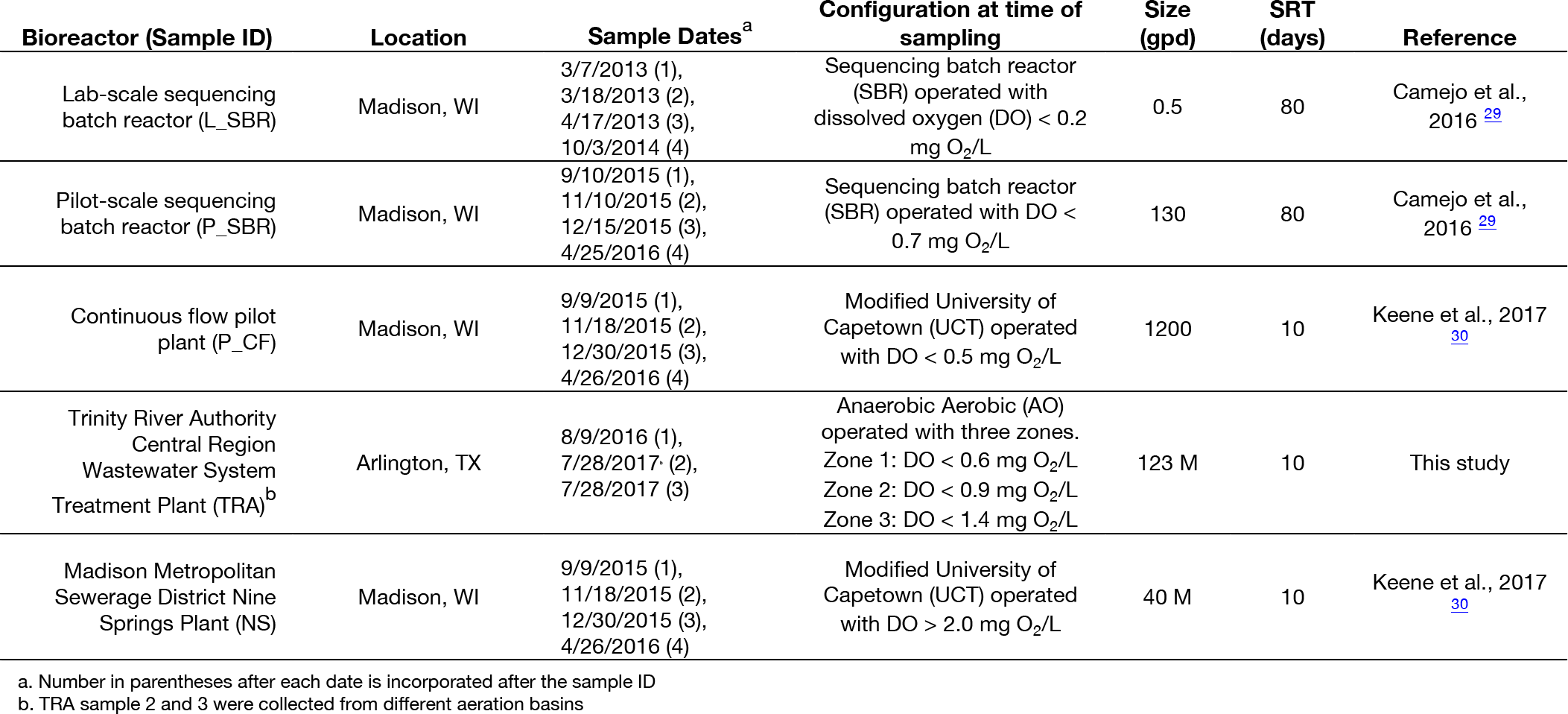
Bioreactor sample characteristics

| Bioreactor (Sample ID) | Location | Sample Dates<sup>a</sup> | Configuration at time of sampling | Size (gpd) | SRT (days) | Reference |
| --- | --- | --- | --- | --- | --- | --- |
| Lab-scale sequencing batch reactor (L_SBR) | Madison, WI | 3/7/2013 (1),<br>3/18/2013 (2),<br>4/17/2013 (3),<br>10/3/2014 (4) | Sequencing batch reactor (SBR) operated with dissolved oxygen (DO) < 0.2 mg O<sub>2</sub>/L | 0.5 | 80 | Camejo et al., 2016 29 |
| Pilot-scale sequencing batch reactor (P_SBR) | Madison, WI | 9/10/2015 (1),<br>11/10/2015 (2),<br>12/15/2015 (3),<br>4/25/2016 (4) | Sequencing batch reactor (SBR) operated with DO < 0.7 mg O<sub>2</sub>/L | 130 | 80 | Camejo et al., 2016 29 |
| Continuous flow pilot plant (P_CF) | Madison, WI | 9/9/2015 (1),<br>11/18/2015 (2),<br>12/30/2015 (3),<br>4/26/2016 (4) | Modified University of Capetown (UCT) operated with DO < 0.5 mg O<sub>2</sub>/L | 1200 | 10 | Keene et al., 2017 30 |
| Trinity River Authority Central Region Wastewater System Treatment Plant (TRA)<sup>b</sup> | Arlington, TX | 8/9/2016 (1),<br>7/28/2017 (2),<br>7/28/2017 (3) | Anaerobic Aerobic (AO) operated with three zones.<br>Zone 1: DO < 0.6 mg O<sub>2</sub>/L<br>Zone 2: DO < 0.9 mg O<sub>2</sub>/L<br>Zone 3: DO < 1.4 mg O<sub>2</sub>/L | 123 M | 10 | This study |
| Madison Metropolitan Sewerage District Nine Springs Plant (NS) | Madison, WI | 9/9/2015 (1),<br>11/18/2015 (2),<br>12/30/2015 (3),<br>4/26/2016 (4) | Modified University of Capetown (UCT) operated with DO > 2.0 mg O<sub>2</sub>/L | 40 M | 10 | Keene et al., 2017 30 |

a. Number in parentheses after each date is incorporated after the sample ID
b. TRA sample 2 and 3 were collected from different aeration basins

### Design of primers to detect comammox ammonia monooxygenase gene *amoA*

A collection of full-length *amoA* and full-length particulate methane monooxygenase (*pmoA*) gene sequences, obtained from the National Center for Biotechnology Information (NCBI) GenBank database,^18^ were used for primer design. All *amoA* and *pmoA* sequences were aligned using the ‘AlignSeqs’ command in the DECIPHER “R” package.^19, 20^ This aligned database was then submitted to DECIPHER’s Design Primers web tool.^21^ Sequences corresponding to *Ca.* N. nitrosa, *Ca.* N. inopinata, and *Ca.* N. nitrificans were individually selected as target groups for primer design, which used the following parameters: primer length ranging from 17 – 26 nucleotides with up to 1 permutation, PCR product amplicon length of 100 – 450 bp, 100% target group coverage, and Taq 3’-end Model to improve specificity. The primer design tool also used the same reaction conditions in all cases: [Na^+^] 70 mM, [Mg_2_^+^] 3 mM, [dNTPs] 0.8 mM, annealing temperature of 64°C, [primers] 400 nM. Amplification products were verified by agarose (2%) gel electrophoresis.

### Quantitative real-time polymerase chain reaction (qPCR)

Quantification of total 16S rRNA genes, total comammox *amoA* genes, clades A and B comammox *amoA*, as well as Ca. N. nitrosa, *Ca.* N. inopinata, and *Ca.* N. nitrificans *amoA* in each DNA sample was carried out by qPCR. All qPCR assays were performed on a Roche LightCycler® 480 high-throughput real-time PCR system using white LightCycler® 480 multiwell plates and the associated LightCycler® 480 sealing foils (Roche Molecular Systems, Inc., Pleasanton, CA, USA). All environmental DNA samples were diluted to 10 ng/μL. All qPCR assays were prepared in Bio-Rad iQ SYBR Green Supermix (Bio-Rad, Hurcules, CA, USA), containing 50 U/mL iTaq DNA polymerase, 1.6 mM dNTPs, 100 mM KCl, 40 mM Tris-HCl, 6 mM MgCl_2_, 20 nM fluorescein, and stabilizers (10 μL per reaction). Nuclease free water (4.4 μL), environmental DNA or standard (4 μL), and PCR primers (0.8 μL of 10 mM each primer) were then added to each reaction for a final volume of 20 μL per reaction. Triplicate reactions were prepared for each sample.

Amplification of total comammox *amoA* was performed according to Fowler et al.^7^ Amplification of clade A and clade B comammox *amoA* was performed using the equimolar primer mixtures according to Pjevac et al.^17^ Amplification of total 16S rRNA was performed using the 16S rRNA-targeted primer pair 341f/785r according to Thijs et al.^22^ (Table 2). Methods for standard preparation are provided in the supporting information.

**Table 2.**
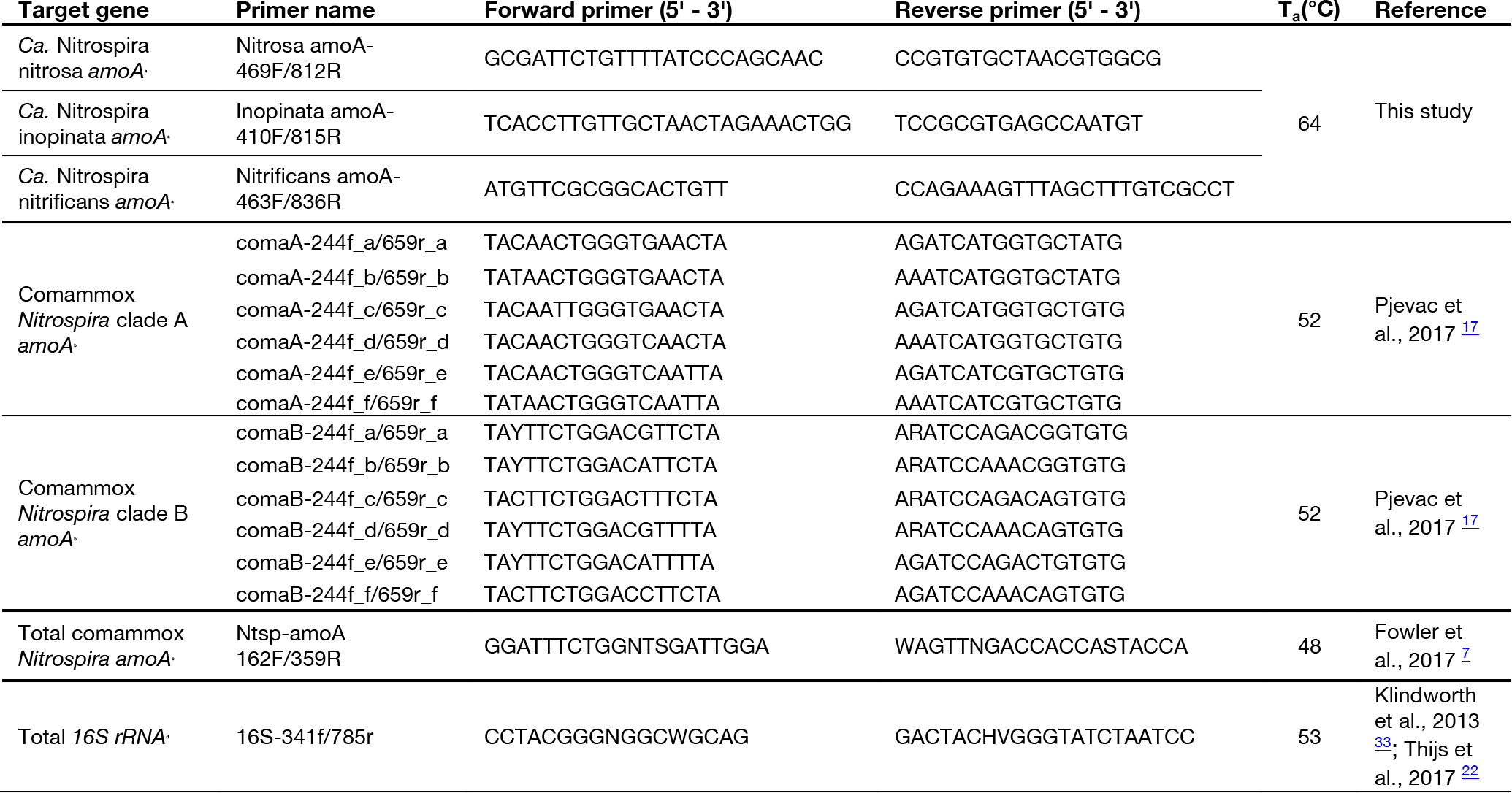
Primers used for qPCR

The thermal cycling protocol for qPCR using the novel *amoA* primers designed in this study (Table 2) was as follows: initial denaturation step at 95 °C for 10 min, followed by 45 cycles of initial denaturation at 95 °C for 10 s, annealing at 64 °C for 30 s, and extension at 72 °C for 30 s. Fluorescence was measured at 72 °C for amplicon quantification. After amplification, an amplicon melting curve was recorded in 0.25 °C steps between 65 and 97°C. Melting peaks were obtained by plotting the negative first derivative of fluorescence against temperature. Although 30 cycles is typically sufficient for quantification of targets in qPCR, the thermal cycling was extended to evaluate potential non-specific amplification with the newly designed primers.^21^

## Results

### Total comammox and clade-level comammox detection

For broad comammox *amoA* quantification, three published primer sets were compared using standards and nineteen environmental samples originating from five bioreactors (Table 1). The Ntsp-amoA 162F/359R primer set,^7^ designed to target total comammox *Nitrospira amoA* (Table 2), amplifies a 198 bp fragment at primer binding regions between positions 162-182 and 339-359 bp (Figure 1). The comaA-244f/659R and comaB-244f/659r primer sets,^17^ designed to differentiate between clades A and B of comammox (Table 2), amplify a 415 bp fragment at *amoA* primer binding regions between positions 244-261 and 643-659 bp (Figure 1). The standard curves for these assays were linear over a minimum of five orders of magnitude and had high coefficients of determination (R^2^ > 0.990); although linear range, amplification efficiency, and limit of detection and quantification varied depending on the standard used (Table 3; Table S1). The three standards used also revealed that the amplified *amoA* of comammox have a wide range of GC content, from 49% to 58%, which resulted in melting temperatures (T_m_) ranging between 84.4 °C and 88.9 °C (Table 3).

**Figure 1.**
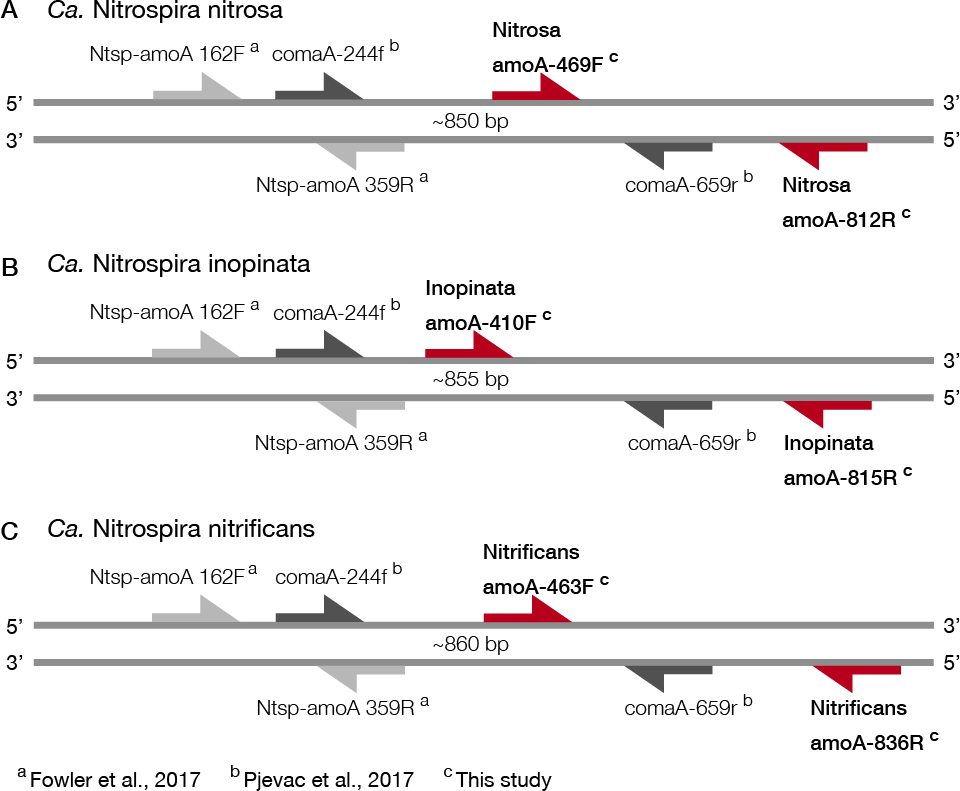
Visual representation of primer amplification regions with A) *Ca.* N. nitrosa *amoA*, B) *Ca.* N. inopinata *amoA*, and C) *Ca.* N. nitrificans *amoA*. The new primers, labeled with red arrows, were designed using full-length *amoA* sequences and novel primer binding region was discovered near the end of each *amoA* gene (approximately between 400 and 800 bp). Base pair locations are indicated in the primer names. Most publicly available published *amoA* sequences are partial length and are located between approximately 244 bp and 659 bp. Amplification with clade A-specific equimolar primer mixtures occur in this region, while amplification with the total comammox primers occur between 162 bp and 359 bp.

**Table 3.** qPCR performance with three distinct comammox *amoA* standard templates

| | Primer set | Amplicon size (bp) | Amplicon GC content (%) | Average $T_m$ (°C) | PCR efficiency (%) | Limit of Detection (copies/ng DNA) | Limit of Quantification (copies/ng DNA) |
| --- | --- | --- | --- | --- | --- | --- | --- |
| Comammox *Nitrospira* clade A | **Standard template: *Ca. N. nitrosa amoA*** |  |  |  |  |  |  |
|  | Nitrosa amoA-469F/812R<sup>a</sup> | 344 | 57 | 88.8 ± 0.08 | 96 ± 3.0 | 7 | 9 |
|  | Ntsp-amoA 162F/359R<sup>b</sup> | 198 | 54 | 86.1 ± 0.08 | 91 ± 2.6 | 30 | 95 |
|  | comaA-244f/659r<sup>c</sup> | 415 | 58 | 88.9 ± 0.09 | 95 ± 2.7 | 6 | 13 |
|  | **Standard template: *Ca. N. inopinata amoA*** |  |  |  |  |  |  |
|  | Inopinata amoA-410F/815R<sup>a</sup> | 406 | 49 | 85.6 ± 0.05 | 90 ± 3.0 | 1 | 1 |
|  | Ntsp-amoA 162F/349R<sup>b</sup> | 198 | 49 | 84.4 ± 0.60 | 82 ± 3.4 | 29 | 37 |
|  | comaA-244f/659r<sup>c</sup> | 415 | 50 | 86.0 ± 0.10 | 86 ± 2.5 | 35 | 43 |
|  | **Standard template: *Ca. N. nitrificans amoA*** |  |  |  |  |  |  |
|  | Nitrificans amoA-463F/836R<sup>a</sup> | 374 | 52 | 86.7 ± 0.10 | 92 ± 2.4 | 1 | 3 |
|  | Ntsp-amoA 162F/359R<sup>b</sup> | 198 | 57 | 87.3 ± 0.27 | 96 ± 1.8 | 6 | 8 |
|  | comaA-244f/659r<sup>c</sup> | 415 | 56 | 88.2 ± 0.07 | 93 ± 2.9 | 5 | 6 |

a. This study
b. Fowler et al., 2017
c. Pjevac et al., 2017

In the total comammox assay, the melting curves for the standards were narrow and unimodal above ~10^1^ − 10^2^ copies (Figure S1 (A-C)). However, melting curves from the environmental samples (Figure S1 (D-H)) and the agarose gel of qPCR products (Figure S2) showed unspecific amplifications resulting in three false positive detections (amplification did not correspond to expected product size) and overestimation of comammox abundance (multiple products seen, including one with the expected product size) in eight samples (Figure 2A).

**Figure 2.**
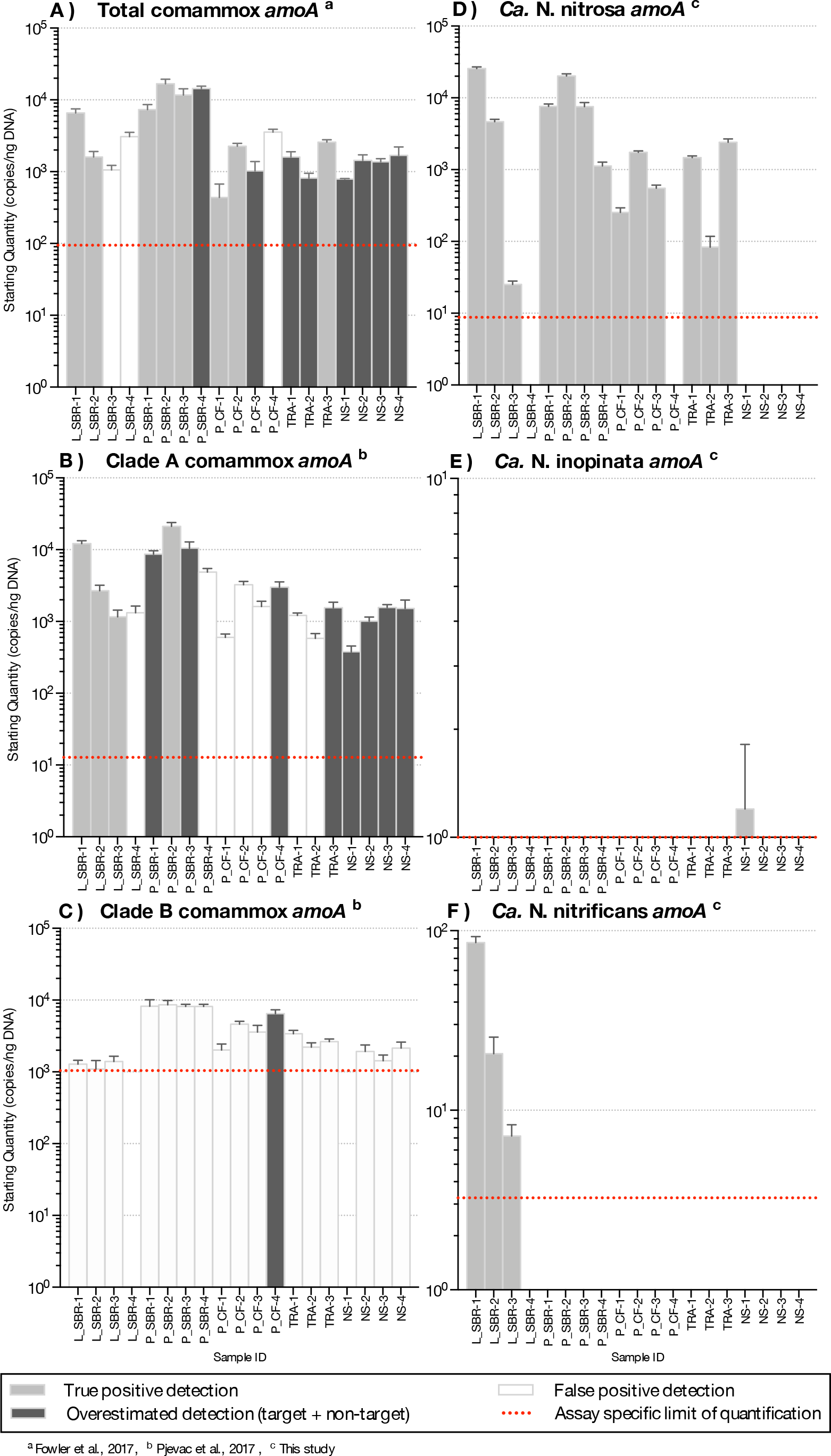
Abundance of *amoA* genes (gene copies/ng DNA) on a log-scale from comammox *Nitrospira* using the (A) clade A primers, (B) clade B primers (C) total comammox primers, (D) the newly designed *Ca.* N. nitrosa specific primers (E) *Ca.* N. inopinata specific primers and (F) *Ca.* N. nitrificans specific primers from various time series samples originating from five bioreactors. LS, lab-scale sequencing batch reactor; P_SBR, pilot-scale sequencing batch reactor; P_CF, continuous flow pilot plant; TRA, Trinity River WWTP; NS, Nine Springs WWTP. Error bars show the standard deviation of the triplicate samples. The color of each column indicates whether there was a true positive detection, an overestimated detection, or a false positive detection. Detections below the limit of quantification are excluded. Note the change in y-axis limits for (E) N. inopinata specific primers and (F) N. nitrificans specific primers.

In the clade A comammox assay, all standards with approximately 10^2^ *amoA* copies or less presented two melting peaks, the expected target peak at approximately 88.9 °C, 86.0 °C, and 88.2 °C for *Ca.* N. nitrosa, *Ca.* N. inopinata, and *Ca.* N. nitrificans *amoA*, respectively, plus a sizeable off-target peak at approximately 70 °C (Figure S3 (A-C)). Higher copy numbers produced narrow, unimodal melting peaks at the expected T_m_. The agarose gel performed with the qPCR products showed a single band for the standards; however, eight of the nineteen environmental samples contained multiple amplified products including a potential target amplicon (Figure S4) leading to overestimated gene abundance (Figure 2B). Amplification of off-target products in seven additional samples produced artifact-associated fluorescence that was well over the limit of quantification and translated into false positive predictions of the same order of magnitude as positive samples (Figure 2B).

Clade B standards with approximately 10^3^ *amoA* copies or less also contained two melting peaks, an off-target peak at approximately 72 °C and the expected target peak at approximately 89 °C (Figure S5 (A)). In the clade B assay, fifteen samples had a target melting peak at the expected T_m_, however, the agarose gel revealed that these samples had abundant off-target products that were approximately 300 bp long— smaller than the expected 415 bp target size (Figure S6). Only one sample (P_CF-4) appeared to contain an amplicon of the expected size (Figure S6), but also contained several other off-target amplicons. Thus, none of the samples tested were confirmed to be positive for presence of clade B comammox.

Overall, the presence, frequency, and influence of unspecific amplification in the total comammox^7^ assay and in the clade A and clade B^17^ comammox assays were significant and did not allow for accurate detection and quantification of comammox *Nitrospira* in the environmental samples tested (Figure 2 (A-C)). Additionally, we found that the GC content variability of the targeted products from the total comammox^7^ and clade A assays^17^ greatly influenced melting characteristics of the known standards included in the tests; thus making it impossible to differentiate between a true positive detection and false positive detection without relying on agarose gel electrophoresis of amplified qPCR products.

### Design of species-specific primer sets

Since the broad range comammox primers did not provide satisfactory results with our environmental samples, we opted to focus our evaluations on the presence of specific comammox species. With this objective in mind, to detect and quantify comammox belonging to the *Candidatus Nitrospira* species currently described in the literature (*Ca.* N. nitrosa, *Ca.* N. inopinata, and *Ca.* N. nitrificans) we designed qPCR primer sets specifically targeting each of these species. For this design, we use a dataset of 27 full-length *amoA* gene sequences from AOB, AOA, and comammox, and 15 full-length *pmoA* gene sequences from methanotrophs. We limited the database to full-length *amoA* and *pmoA* sequences in order to allow for discovery of primer-binding regions outside of the fragments amplified by conventional *amoA* primer sets.^12, 23^ We used the Design Primers option in DECIPHER^21^ with each one of the candidate species as the target group and other *amoA/pmoA* sequences entered as closely related groups that should not be amplified. In addition, for all designs, the annealing temperature and PCR conditions were fixed (see Materials and Methods) so that all three species-level primer sets could be simultaneously used in a single thermocycler run.

For *Ca.* N. nitrosa, the two *amoA* copies identified in the *Ca.* N. nitrosa genome^2^ were used as the target group. The design algorithm identified primer binding regions at positions 469-493 and 794-812, predicting amplification of a 344 bp fragment (Figure 1). The target regions had more than 4 mismatches to all other sequences, predicting a 100% specificity to the target group.

A single sequence^1^ served as the target group for the design of *Ca.* N. inopinata primers, resulting in primer binding sites at 410-436 and 798-815, for a 406 bp fragment amplification. In this primer set, there were more than 5 mismatches to other *amoA* and *pmoA* sequences used as non-targets, also predicting 100% specificity.

For *Ca.* N. nitrificans, two sequences were used as targets; the one in the *Ca.* N. nitrificans genome^2^ and the one from *Nitrospira* sp. Ga0074138^3^, which had 91.4% sequence identity to the *Ca.* N. nitrificans sequence. The primers amplified a 374 bp fragment, between positions 463 and 836. The predicted specificity was again 100%, with more than 4 mismatches to other sequences used in the design dataset.

Since the sequence dataset used for design was small, we searched for additional *amoA* sequences that would contain the target regions of the new primer sets. However, this analysis was limited because most of the published *amoA* sequences contain the target side of the forward primers, but not the site targeted by the reverse primer (Figure 1). In this search, we found 33 nearly full-length sequences that clustered with the comammox sequences and contained the target sites for both primers (Figure 3), including *amoA* sequences from comammox metagenomes that have been recently described in the literature.^1–3, 16, 17, 24^

**Figure 3.**
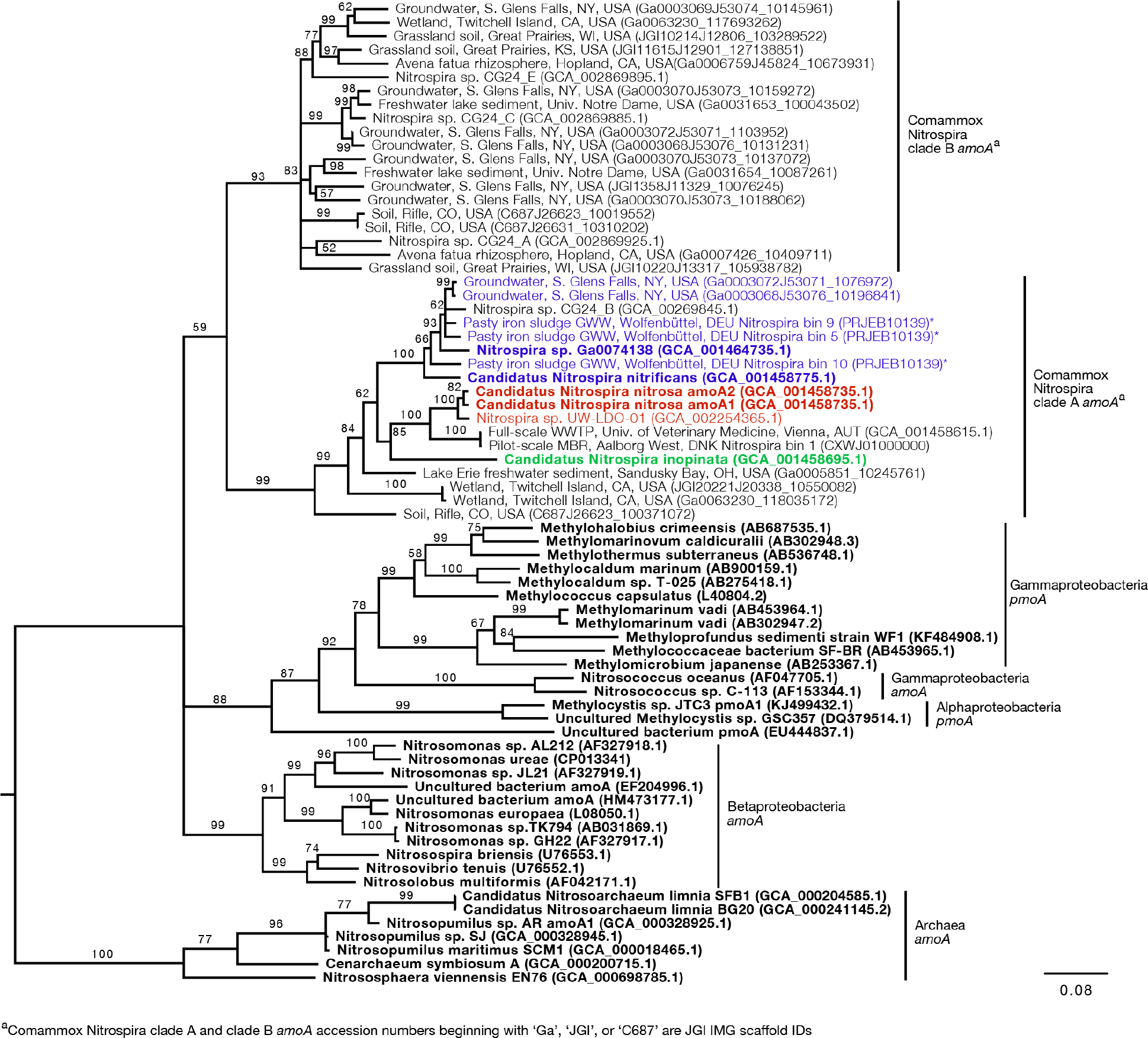
Neighbor-joining consensus tree generated from an alignment of full-length and near full-length *amoA* and *pmoA* gene sequences (> 600bp), rooting with archaea *amoA*. Bootstrap values, shown at the nodes where the value was greater than 50, are based on 10,000 trials. Bold sequence names were included in the new primer design. Blue, red, and green text are amplified by the *Ca.* N. nitrificans, *Ca.* N. nitrosa, and *Ca.* N. inopinata primer sets, respectively. Asterisks indicate that the gene is amplified but with a reduced efficiency due to base pair mismatches. The scale bar indicates the number of nucleotide substitutions per site. Accession numbers are presented after the sequence names. Acronyms were used for groundwater well (GWW), wastewater treatment plant (WWTP), and membrane bioreactor (MBR).

An *in silico* primer-target analysis (Figure 4) shows that *N. sp.* UW-LDO-01^16^, originally described as a strain of *Ca.* N. nitrosa, has perfect matches to the newly designed primers targeting this species. Two *amoA* sequences (Full scale WWTP, Univ. of Veterinary Medicine, Vienna, AUT, GCA_001458615.1; Pilot-Scale MBR, Aalborg West, DNK Nitrospira bin 1, CXWJ01000000) clustered together and near the *Ca.* N. nitrosa cluster (Figure 3). The average nucleotide identity (ANI) of their genomes compared to *Ca.* N. nitrosa is 84.5% and 85%, respectively (Figure S7), lower than the typical cuttoff for species definition (ANI > 94%).^25^ In agreement, we predict that their *amoA* will not be amplified by the nitrosa-specific primer set due to multiple mismatches with forward and reverse primers (Figure 4). The *amoA* sequence of *Nitrospira sp.* CG24_B^*24*^ clustered with *Ca.* N. nitrificans (Figure 3). However, the ANI of these two genomes (86.3%; Figure S7) is below the typical cutoff for species definition and the *in silico* analysis predicts that *Nitrospira sp.* CG24_B *amoA* will not be amplified with the nitrificans-specific primers (Figure 4). Three additional *amoA* sequences clustering with *Ca.* N. nitrificans include those from the draft genomes of ‘Pasty iron sludge GWW N. bin 5, N. bin 9, and N. bin 10 (PRJEB10139)’^1^ (Figure 3), which contain fewer mismatches (Figure 4) and are predicted to partially amplify (below 56% effiency) with the *Ca.* N. nitrificans primers despite ANI below the species cutoff (Figure S7). Overall, the *amoA* phylogeny close to *Ca.* N. nitrificans remains unresolved, and therefore, the designed primer set for this species will likely require future refinement.

**Figure 4.**
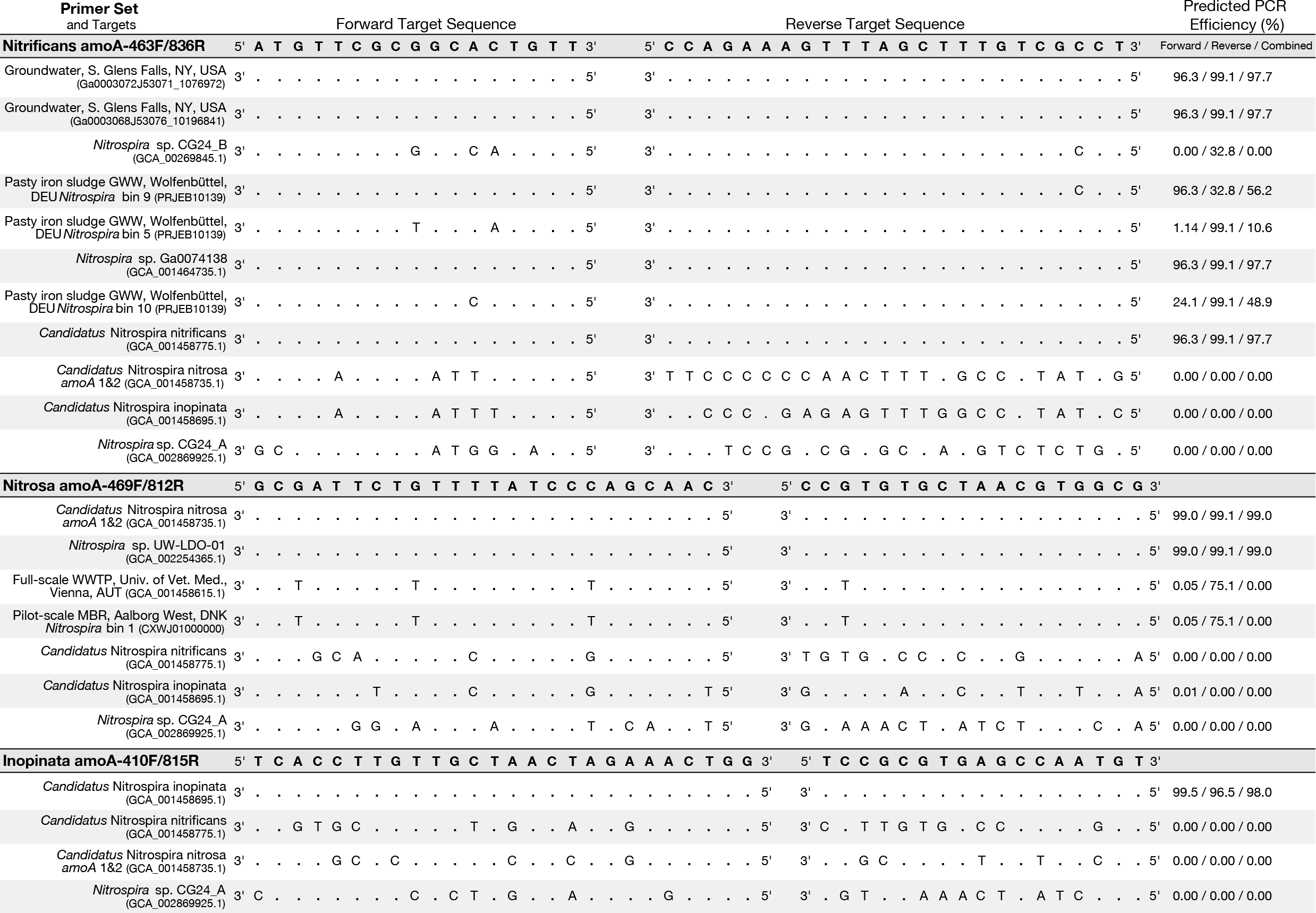
Primer-target mismatch analysis for the newly designed primers and near full-length comammox clade A *amoA* greater than 600 bp. Predicted PCR efficiency is reported separately for the forward and reverse primers in addition to a combined amplification efficiency.

Sequences from clade B comammox were not included in dataset used for primer design, and therefore, an in silico analysis was also performed with these sequences (Figure 4 shows *N. sp*. CG24_A *amoA* as an example of clade B comammox). In all cases, the sequences had greater than 4 mismatches and are predicted to not amplify with any of the new species-specific primer sets.

The standard curves for all three primer sets were linear (correlation coefficient R^2^ ≥ 0.996) over 7 orders of magnitude (Table S1). The amplification efficiency for *Ca.* N. nitrosa, *Ca.* N. inopinata, and *Ca.* N. nitrificans *amoA* was 96 ± 3.0%, 90 ± 3.0%, and 92 ± 2.4%, respectively (Table 3). Melting curve analysis of all standards showed amplification of the target product without primer dimer artifacts, represented by a strong fluorescence signal producing a single melting peak at approximately 88.8 °C for *Ca.* N. nitrosa (Figure S8 (A)), 85.6 °C for *Ca.* N. inopinata, and 86.7 °C for *Ca.* N. nitrificans (Figure S9 (A) – S10 (A)).

### Environmental detection of candidate comammox species

Using the new species-specific primer sets developed in this study (Nitrosa amoA-469f/812r, Inopinata amoA-410f/815r, and Nitrificans amoA-463f/836r), we evaluated the samples originating from the low DO BNR bioreactors (Table 1). With the new primers, positive detections obtained with melting curve analysis (Figure S8-S10) correlated well with a positive target amplicon in the agarose gel (Figure S10-S11). Additionally, cross-hybridization and primer dimers were not observed following PCR with agarose gel electrophoresis with the standard templates (Figure S12).

Comammox *amoA* belonging to *Ca.* N. nitrosa were detected in all environmental samples. Agarose gel electrophoresis of the qPCR amplified products validated these results, since a single amplicon with the expected length (344-bp) was obtained from all 19 samples (Figure S11). Visible products with the expected length were also present for two samples where the predicted abundance was below the limit of detection (L_SBR-4, and NS-4) and four samples below the limit of quantification (P_CF-4, NS-1, NS-2, and NS-3) (Table 3, Figure S11), indicating a high level of sensitivity. The melting curve analysis (Figure S8) confirmed the specificity of the amplifications. *Ca.* N. nitrosa abundance was greater than 10^3^ copies *amoA*/ng DNA in nine samples originating from low DO bioreactors L_SBR, P_SBR, P_CF, and TRA (Figure 2D). The maximum number of *amoA* copies was obtained from L_SBR-1, with approximately 2.5 × 10^4^ copies *amoA*/ng DNA (Figure 2D).

Compared to the quantification of total comammox^7^, *Ca.* N. nitrosa comprised an average of 340 ± 70%, 96 ± 29%, 67 ± 13%, and 93% of the total comammox population in the true positive detections obtained from L_SBR, P_SBR, P_CF, and TRA, respectively (Figure 2A, 2D). These results indicate that the total comammox primers^7^ may have underestimated *Ca.* N. nitrosa species in samples with an abundant comammox population (L_SBR and P_SBR). A closer look at the primer-target region for the total comammox primers^7^ against *Ca.* N. nitrosa *amoA* revealed 2 mismatches in both the forward and reverse primers (Table S2), as mismatches were allowed in the original total comammox primer design.^7^ Moreover, we observed reduced total comammox^7^ efficiency when *Ca.* N. nitrosa *amoA* was present at less than 10^3^ copies (Table S1). Thus, total comammox^7^ primer mismatches to *Ca.* N. nitrosa *amoA* may contribute to an overall reduced amplification efficiency and underestimation of this species in environmental samples that contain a true positive detection. All in all, the new species-specific primers show that *Ca.* N. nitrosa is an important member of the comammox population in wastewater treatment plants and may contribute to low oxygen nitrification (as seen in L_SBR, P_SBR, P_CF and TRA and below the limit of quantification in NS).

Comammox *amoA* belonging to *Ca.* N. inopinata and *Ca.* N. nitrificans were less frequent and were only detected with less than 10^2^ copies *amoA*/ng DNA in all samples (Figure 2E, 2F; Figure S13). N. inopinata *amoA* was only detected in one sample from the Madison Metropolitan Sewerage District Nine Springs plant (NS), NS-1, with just 1 copy *amoA*/ng DNA (Figure 2E). *Ca.* N. nitrificans was only detected above the limit of quantification in L_SBR-1, L_SBR-2, and L_SBR-3 with approximately 86, 21, and 7 copies *amoA*/ng DNA, respectively (Figure 2F).

Overall, comammox *amoA* belonging to *Ca.* N. inopinata or *Ca.* N. nitrificans were minor contributors to the total bacteria population after normalization (with relative abundances less than 0.03%) (Figure S14).

## Discussion

Since the late 1800s, nitrification had been described as a division of labor between two phylogenetically distinct and specialized chemolithoautotrophs, the ammonia oxidizers and the nitrite oxidizers.^26^ This perception was not challenged for over a century, until Costa et al.,^27^ hypothesized that a single nitrifying bacterium combining ammonia oxidation and nitrite oxidation should exist in nature.^27^ Nearly one decade later, in late 2015, two separate research groups discovered, cultivated, and characterized the ‘missing’ comammox organism from an aquaculture system and a deep oil exploration well, respectively.^1, 2^ To date, there is still uncertainty about the occurrence of the novel *Nitrospira-like* comammox organisms in full scale activated sludge systems. While some studies have suggested that comammox are not relevant to conventional wastewater treatment^4, 28^, no studies have been published that specifically attempt to quantify comammox in wastewater treatment plants with low DO. Since comammox were discovered in low oxygen environments^1, 2^ they may be able to efficiently utilize oxygen and may be important to low DO BNR.

Long-term low DO total nitrogen removal has been successfully demonstrated in laboratory^29^ and pilot-scale^29, 30^ bioreactors seeded with Nine Springs WWTP activated sludge and operated with DO concentrations less than 0.60 mg O_2_/L. Samples analyzed from the Nine Springs WWTP showed presence of *Ca.* N. nitrosa, but below the limit of quantification. The sludge from this WWTP was the seed for the L_SBR, P_SBR, and P_CF reactors, all of which showed presence of *Ca.* N. nitrosa, suggesting that the low DO conditions favored accumulation of *Ca.* N. nitrosa as a potential participant in ammonia oxidation in low DO BNR. Interestingly, *Ca.* N. nitrosa was also abundant in the full-scale TRA system, which is operated with low DO conditions (Table 1), providing independent support for the hypothesis that *Ca.* N. nitrosa is an important contributor to ammonia oxidation in WWTP operated with low DO. *Ca.* N. nitrificans was detected above the limit of quantification in the L_SBR reactor only, whereas Ca. N. inopinata was not detected in any of the low DO reactors, and therefore, out of the three candidate species, *Ca.* N. nitrosa appears to be the only one that becomes enriched under low DO conditions.

When compared to quantification of the total bacterial population by 16S rRNA-targeted qPCR, the greatest relative abundance of *Ca.* N. nitrosa was 10% and 6% of total bacteria in samples originating from the laboratory-scale and pilot-scale sequencing batch reactors, respectively (Figure S14). Although these two reactors were seeded with the same sludge, they were fed from completely different sources (synthetic media for L_SBR and full-scale primary effluent for P_SBR), but operated with low oxygen and a long solids retention time (SRT, both 80 days).^29^ The continuous flow reactors (P_CF and TRA) had lower relative abundances of Ca. *N. nitrosa*, and although operated with low DO, they had much lower SRT (both 10 days; Table 1). Thus, in addition to oxygen, SRT may be a factor that contributes to comammox abundance in low DO reactors.

Detection and quantification of microorganisms via real-time PCR relies on designing primers with good specificity to the targeted organisms, good coverage of the targeted group, and good quality of the experimental results. With any newly discovered target group, the quality of primer sets to achieve accurate quantification depends on the quality of the databases used for design and the design considerations. As more sequences of the targeted organisms become available, designs can only improve. The first primer design for comammox quantification described sets for targeting clade A and B *amoA* within the *Nitrospira* genus^17^, and subsequently another set of primers was published, aiming for greater coverage to target all comammox *amoA*^7^. Our evaluation of these primer sets in environmental samples from low DO BNR plants showed challenges with primer dimer formation when target sequences were not abundant (clade A and clade B primers^17^), unspecific amplifications in environmental samples (all sets), and underestimation with some strains due to primer-target mismatches (total comammox^7^). These challenges are more important when designing for broad target groups. In general, melting curve analyses of qPCR results are helpful for detection of nonspecific PCR products since different fragments will typically appear as distinct melting peaks.^31, 32^ However, the broad detection and quantification of comammox *amoA* has high GC content variability among the comammox organisms described thus far (Table 3). Therefore, a wide range of amplicon melting temperatures are expected when using the total comammox^7^ or clade-level^17^ assays, making melting curve analyses difficult to interpret. We found that verifying amplified products with the total comammox^7^ or clade-level^17^ primers cannot be completed with melting curve analysis alone and will require a subsequent agarose gel electrophoresis of all environmental samples, which is not ideal for routine qPCR applications.

Setting the challenge of broad comammox quantification aside, we aimed at designing primer sets specific to the three candidate species described thus far. Evidently, there is a very small number of sequences representative of these candidate species; therefore, the designed primers inherently have 100% coverage. Specificity depends on finding sufficient differences between target and non-targets sequences^21^, and the results showed this was possible for the three candidate species targeted. As more sequences become available from metagenome assemblies, the specificity of these primers can be re-evaluated. The beginning of such activity was performed in this study (Figure 4). Importantly, the qPCR experiments with the newly designed primers were conducted with a longer number of thermal cycles than typical (45 cycles) to increase the chances of detecting potential non-specific amplifications. This extended thermal cycling confirmed that that the primers are highly specific and are not producing unwanted non-specific amplifications.

Taken together, the species-specific comammox primers designed in this study enabled an analysis of comammox abundance focused solely on the candidate comammox species currently described in the literature. Although a narrowly focused analysis by design, it eliminated the
problems of unspecific amplifications that compromised the use of broad comammox primers, and resulted in strong experimental evidence in support of *Ca.* N. nitrosa being a comammox organism with an important contribution to ammonia oxidation in energy-efficient BNR systems operated under low DO conditions.

## Supporting Information

Detailed information regarding standard preparation, phylogenetic tree construction, and average nucleotide identity calculation are presented. Additional tables describing standard curves, mismatches, and supporting figures of qPCR product agarose gel electrophoresis and qPCR melting curves for all assays are also located in the associated supporting information PDF.

## Acknowledgements

This work was partially supported by funding from the National Science Foundation (CBET-1435661) and the Madison Metropolitan Sewerage District (Madison, WI). We thank Colin Fitzgerald and Leon Downing (Jacobs Engineering Group) for providing collaboration and coordination with the Trinity River Authority (Arlington, TX). We thank Pamela Camejo for providing training for primer design and cloning. We also thank Rahim Ansari, Jonnathan Garcia-Huerta, and Tara Hawes for support with cloning, DNA extraction, and PCR. Finally, we would like to recognize Jackie Bastyr-Cooper for her assistance in the lab with training, protocols and equipment.

